# Metal sensing properties of the disordered loop from the Arabidopsis metal transceptor IRT1

**DOI:** 10.1101/2024.09.03.611018

**Authors:** Virginia Cointry, Reyes Ródenas, Nelly Morellet, Valérie Cotelle, Julie Neveu, Grégory Vert

## Abstract

The plant IRT1 iron transporter is a plasma membrane protein that takes up iron in root upon iron-limited conditions. Besides its primary metal substrate iron, IRT1 also transports other divalent metals that overaccumulate in plants when soil iron is low and *IRT1* is highly expressed. We previously reported that the intracellular regulatory loop between transmembrane helices TM4 and TM5, comprising IRT1 residues from 144 to 185, is involved in the post-translational regulation of IRT1 by its non-iron metal substrates. Upon excess of zinc, IRT1 (144-185) undergoes phosphorylation by the CIPK23 kinase followed by its ubiquitination by IDF1 to target IRT1 for vacuolar degradation. This zinc-dependent downregulation of IRT1 requires the presence of four histidine (H) residues in IRT1 loop, that directly bind zinc. However, how selective metal binding is achieved and how this allows downstream regulation to take place is largely unknown. Here, we characterized the metal binding properties and structure of IRT1 loop to better understand the molecular basis of non-iron metal sensing and signaling. Using a combination of circular dichroism and NMR, we demonstrate that zinc and manganese bind to IRT1 loop with nanomolar range affinity, and that metal binding does not trigger structuration of the loop. We prove that zinc and manganese binding is mediated by the four H residues and identify aspartic acid (D) residue D173 as helping in metal coordination and participating to metal sensing and metal-dependent degradation of IRT1 in plants. Altogether, our data provide further evidence of how the regulatory loop of IRT1 senses high cytosolic divalent metal concentrations to regulate metal uptake in plants.

## Introduction

The first-row transition metal iron is the most abundant and used metal in cells (1). Iron is widely used as a cofactor, participating in oxidation-reduction reactions. In plants, iron is of utmost importance due to its involvement in photosynthesis electron transfer reactions (2). In nature, iron is often unavailable to organisms including plants because it is found as insoluble ferric oxide/hydroxide. Lack of iron deeply impacts plant growth and leads to severe leaf chlorosis (2). On the other hand, an excess of iron is detrimental to cells due to its redox properties and participation in the Fenton reaction that leads to reactive oxygen species formation (3). For these reasons, living organisms and plants in particular need to tightly control metal uptake.

Dicotyledonous plants, such as the model plant *Arabidopsis thaliana*, use a three-step iron uptake process based on the acidification-based solubilization of soil ferric iron (Fe^3+^) insoluble oxides and the reduction into ferrous iron (Fe^2+^) prior to transport. This is mediated by the AHA2 proton pump, which secretes proton, and the FRO2 ferric chelate reductase, which reduces Fe^3+^ to Fe^2+^ (4–7). In Arabidopsis, the main transporter involved in the uptake of Fe^2+^ is the Iron-Regulated Transporter 1 (IRT1) (8), one of the founding members of the family of ZIP transporters that are found across all kingdoms of life (9). Arabidopsis IRT1 has a broad substrate range, as it transports other essential divalent heavy metals such as zinc (Zn^2+^), manganese (Mn^2+^), cobalt (Co^2+^) and toxic cadmium (Cd^2+^) ions (8,10–16). *In planta*, IRT1 is the major root iron transporter responsible for iron uptake from the soil under iron-limited conditions. IRT1 protein indeed localizes to the plasma membrane and *IRT1* gene is strongly expressed in iron-deficient root epidermal cells (8,15). Consistently, *irt1-1* knockout mutant plants are highly chlorotic and show growth impairment, a phenotype that can be reversed by the addition of high amounts of iron in watering solutions (8). The chlorosis of *irt1* mutants is however not complemented by the other divalent metal substrates of IRT1, indicating that these constitute secondary substrates that are aspecifically transported.

In recent years, the molecular mechanisms underlying the regulation of *IRT1* expression by its metal substrates have been uncovered. The transcription of *IRT1* is upregulated upon iron starvation through a cascade of bHLH transcription factors (17–23). Most importantly, post-translational regulation of IRT1 by its non-iron metal substrates Zn^2+^, Mn^2+^, Co^2+^ and Cd^2+^ has recently been reported (16,24). Non-iron metal availability indeed regulates the subcellular localization and stability of IRT1. In the absence of non-iron metals, IRT1 sits at the cell surface to take up low available iron from the soil. Increasing non-iron metal levels lead to partial IRT1 internalization in early endosomes, as a result of its multi-monoubiquitination on lysine residues K154 and K179 by a yet to be characterized E3 ubiquitin ligase (15). Higher non-iron metal levels trigger IRT1 phosphorylation by the CIPK23 kinase and the recruitment of the E3 ubiquitin ligase IDF1 (16,24). IDF1 elongates monoubiquitin moieties into K63 polyUbiquitin chains, thus leading to IRT1 targeting to the vacuole for degradation (16,24). Fe^2+^ availability has however no influence on IRT1 protein localization (16,24).

The cytosolic regulatory loop of IRT1 located between transmembrane domains TM4 and TM5, IRT1 (amino acid 144-185), has been shown to bind metals *in vitro* (24). A remarkable feature of IRT1 and related transporters is the presence of a histidine-rich motif at the center of such loop. Such repetition of histidine has been shown to be required for the non-iron metal-dependent endocytosis of IRT1, as an IRT1_4HA_ mutant with the four histidines mutated to alanine fails to be degraded upon non-iron metal excess (24). Metal binding to histidine residues was shown to drive the recruitment of CIPK23 (24). IRT1 is therefore considered as a bifunctional transporter-receptor capable of sensing elevated non-iron metal levels and initiating a signaling cascade culminating in its self-degradation to limit the entry of highly reactive non-iron metals. The intricate mechanism allowing metal binding to IRT1 histidine-rich motif to trigger CIPK23 recruitment is however still unclear but was proposed to involve the folding of IRT1 (a.a.144-185) (25).

To obtain better mechanistic insight into how IRT1 senses non-iron metals and how this mediates the recruitment of CIPK23, we biochemically characterized the metal binding properties and the structure of IRT1 loop (a.a. 144-185) by a combination of circular dichroism (CD), NMR spectroscopy and microscale thermophoresis (MST). We report that the regulatory loop of IRT1 binds Zn^2+^ and Mn^2+^ *in vitro* using the four histidine residues with nanomolar affinities, and we also uncover the role of residue D173 in Zn^2+^ and Mn^2+^ coordination and sensing during IRT1 degradation in response to non-iron metal excess. Furthermore, we show that the regulatory loop of IRT1 is a disordered domain both in the absence and presence of Zn^2+^, despite of experiencing small structural changes in presence of Zn^2+^. Overall, our work provides additional biochemical and structural insight into the ZIP family of metal transporters.

## Results

### The regulatory loop of IRT1 is disordered

Considering that to date eukaryotic ZIP transporters cannot be expressed and purified from heterologous systems, we focused on the regulatory loop of IRT1 that carries important residues for metal sensing and response (24), as previously done for hZIP4 (26,27). To get a first glimpse of the structural features of the regulatory loop of IRT1 (a.a. 144-185), we performed far-UV circular dichroism (CD) on the corresponding peptide chemically synthesized. The ^143^DSMATSLYTSKNAVGIMPHGHGHGHGPANDVTLPIKEDDSSN^186^ peptide sequence encompasses all important residues required for phosphorylation and ubiquitination, and contains the four histidine residues that have been previously associated with metal sensing (16,24). Structure prediction using the PONDR (http://www.pondr.com) software suggests a high degree of disorder in this region (Figure S1), with 45% of the residues being disorder promoting (A, R, G, Q, S, P, E and K), and only 21% being rather order promoting (W, C, F, I, Y, V, L, and N). In accordance with software predictions, CD spectra recorded from 190 to 240 nm at pH 6.7 at room temperature showed a spectrum typical of a peptide in a random coil conformation, with a single negative peak at 198 nm (28) (Figure 1A, gray line, 0 eq).

**Figure 1.**
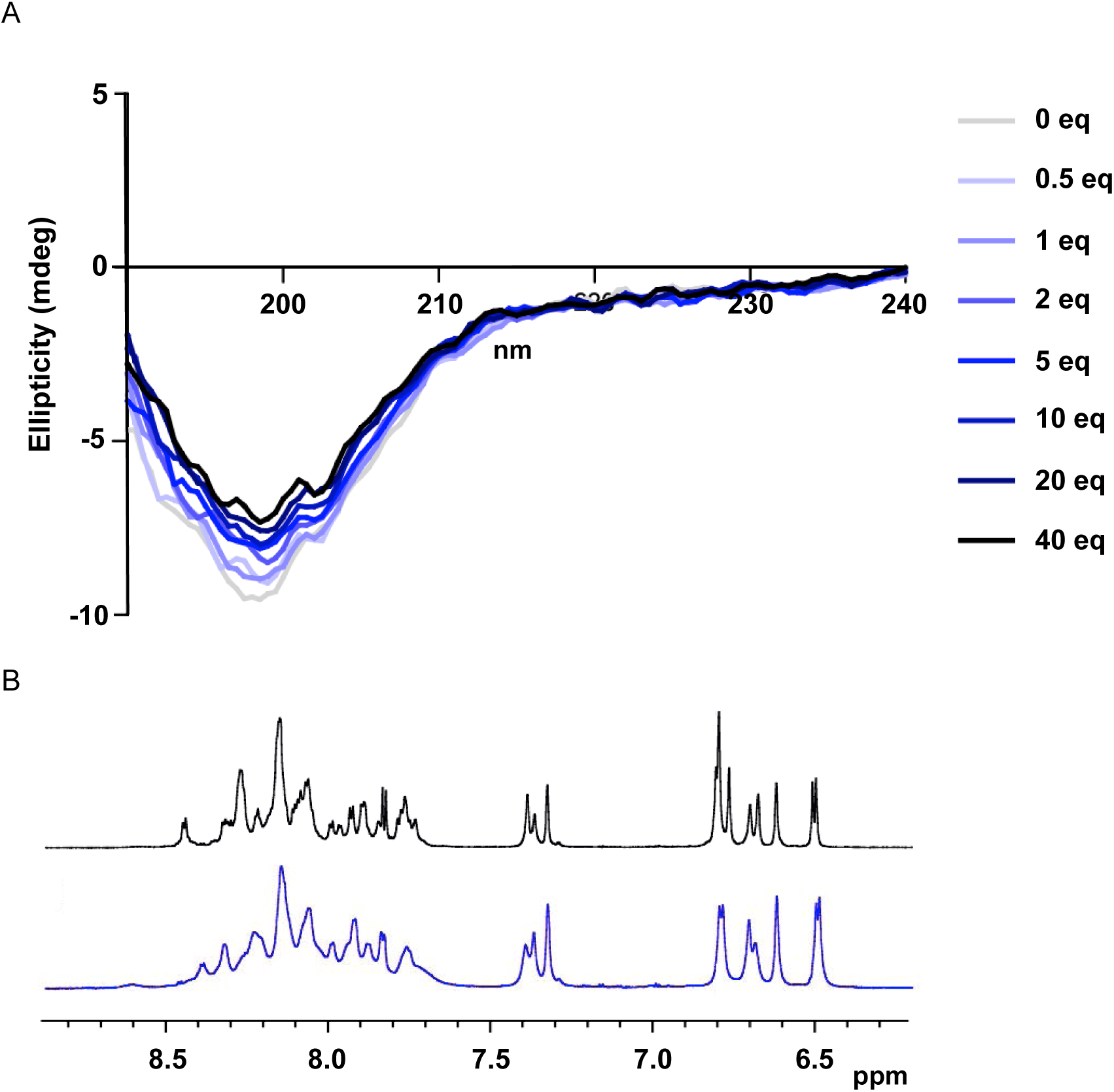
The IRT1 regulatory loop is disordered. A) Far-UV circular dichroism spectra (CD) of IRT1 (144-185). Equivalents of Zn^2+^ (0 to 40 eq) were added to IRT1 (144-185) peptide and CD spectra were recorded from 190 to 240 nm. Corrections of the final peptide concentrations were done with each addition of Zn^2+^. B) One dimension (1D) ^1^H NMR spectra of IRT1 (144-185) recorded in the absence (black) and presence (blue) of Zn^2+^. Only the amide and the aromatic protons are shown.

To obtain deeper insight into the structure of IRT1 loop, we turned to NMR spectroscopy. One dimension (1D) ^1^H NMR spectra were recorded at pH 6.7 in the absence of metals. In line with the results obtained by CD, 1D ^1^H NMR spectra were poorly dispersed, a characteristic of peptides in random coil conformations (Figure 1B, black line). Furthermore, two-dimension (2D) H^1^-H^1^ NOESY spectra of IRT1 loop showed that despite our previous observations of the peptide adopting a rather unstructured conformation, medium-range NOEs existed, notably pointing to the presence of a turn involving the ^155^AVGI^160^ hydrophobic residues (α156/HN159, α157/HN159) (Figure S2A). Besides, we observed that the cross-peak intensities of the histidine-rich domain from IRT1 decreased relative to the other signals of the peptide, resulting probably from chemical exchange (Figure 2A). Despite the lack of a clear secondary structure, we generated structures using the NOE-derived distance restraints with the CYANA software (Figure S2B). As expected, except for the ^155^AVGI^160^ turn (Figure S2C), no secondary structure was observed.

**Figure 2.**
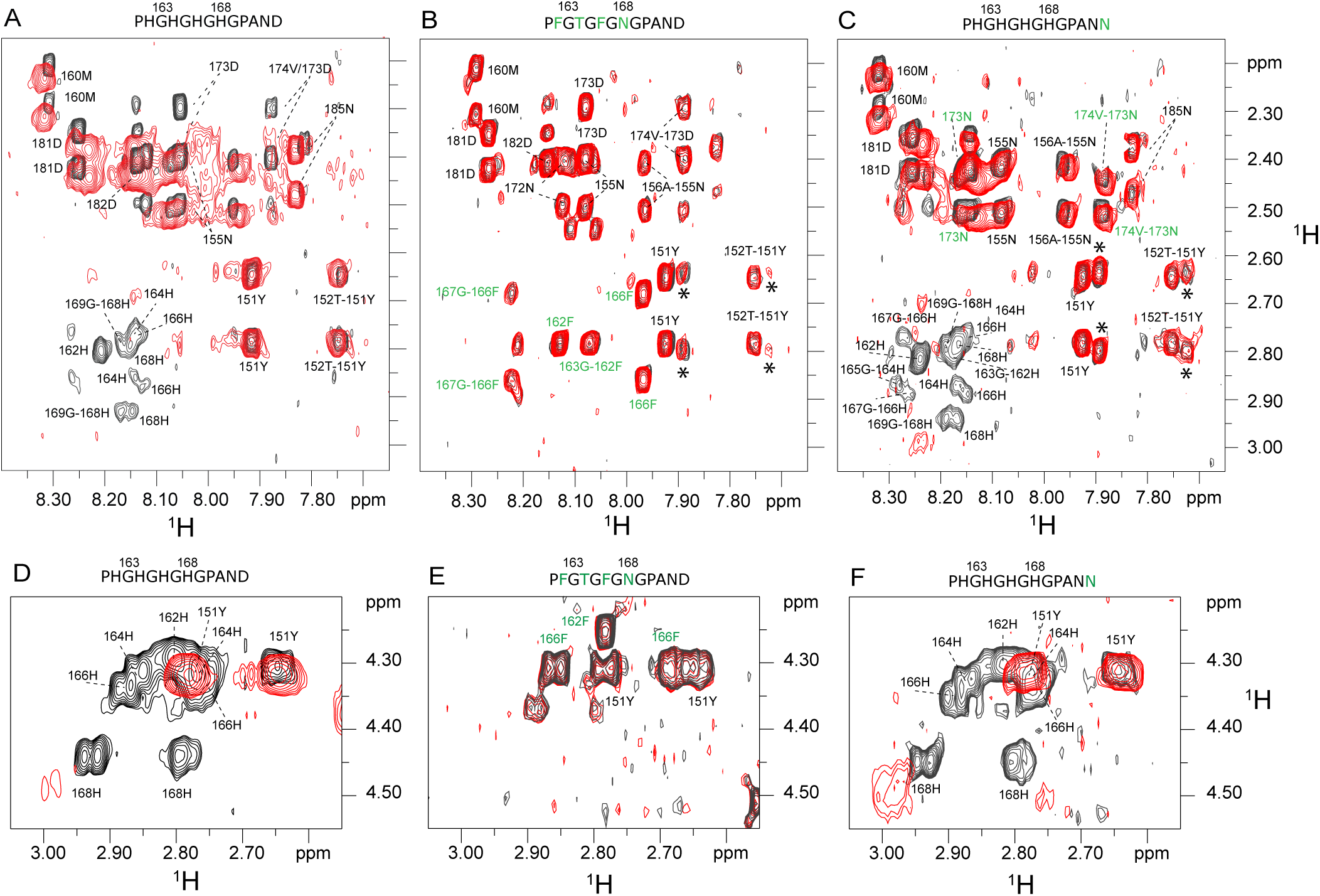
Zinc directly binds to histidine residues in IRT1 regulatory loop. Fingerprint region of the ^1^H-1H NOESY spectrum recorded on IRT1 (144-185) in the presence or absence of Zn^2+^. Several intramolecular and intermolecular NH-CHβ signals are shown for wild-type IRT1 (144-185) (A) and two mutant peptides (B, C); and intramolecular CHα-CHβ signals for wild-type IRT1 (144-185) (D) and two mutant peptides (E, F). The sequences of the domain into which mutations have been introduced are shown above each spectrum and the mutations are highlighted in green. The spectra recorded in the absence of Zn^2+^ are shown in black and those recorded in the presence of two molar equivalents of Zn^2+^ are shown in red. * Indicate impurities in the sample. Annotations with a dash indicate cross-peak between inter residue i.e. NHi-CHβi-1, and without dash cross-peaks between intra residue i.e. HNi-Hβi.

### Impact of metal binding on the structure of IRT1 regulatory loop

In order to determine whether the structural properties of IRT1 loop are impacted by interaction with the secondary metal substrates of IRT1, we performed a titration by sequential equimolar additions of Zn^2+^ on the peptide followed by recording of CD spectra (Figure 1A). With each addition of Zn^2+^, the final concentration of the peptide was corrected to avoid dilution effects. We observed a saturable increase in ellipticity with each addition of the substrate but the overall shape of the spectra remained unchanged. Overall, the binding of Zn^2+^ did not seem to impact the structure of IRT1 loop, as visualized by CD. Similarly, 1D H^1^ NMR spectra in presence of Zn^2+^ showed a similar dispersion of the signals compared to the spectra recorded in absence of metal, again suggesting that even in presence of Zn^2+^ IRT1 loop remains disordered (Figure 1B, blue line). Interestingly, the signals are broadened upon sequential addition of Zn^2+^, indicating that IRT1 loop undergoes small structural changes probably induced by the binding of Zn^2+^ ions (Figure 1A, blue colored lines).

To obtain a deeper understanding of the molecular interaction of IRT1 loop with its substrates, Two-dimension (2D) H^1^-H^1^ NOESY spectra of the IRT1 peptide were compared at pH 6.7 in the absence of metal substrates and in the presence of 2 equivalent of Zn^2+^. Superimposition of these spectra indicated that the histidine-rich motif of IRT1 is implicated in the binding of Zn^2+^ since all the proton resonances of these histidines (H162, H164, H166 and H168) disappeared upon the addition of Zn^2+^ (Figure 2A). This suggests that these residues are involved in the exchange between several conformations in an intermediate regime. Further analysis of our data also indicated that signals corresponding to D173, which is located C-terminal of the histidine-rich stretch, behaved similarly to the histidine residues (Figure 2A). NMR signals corresponding to D173 indeed disappeared after the addition of Zn^2+^, suggesting a possible implication of D173 in metal coordination.

To better characterize the role of individual histidine residues and aspartic acid D173 present in the wild-type IRT1 loop, we decided to also record H^1^-H^1^ NOESY spectra of different mutant peptides. We designed three mutants for the PHGHGHGHGPAND motif where histidine residues were replaced with phenylalanine (F), threonine (T) or asparagine (N) to remove the charges at these positions and maintain closest residue similarity; and a mutant where aspartic acid D173 was replaced by the uncharged asparagine amino acid. We observed that in presence of Zn^2+^, the line width for most of the peptides increased drastically (Figures 2A, C, D, F and S3) except for the mutant peptide with four mutated histidine residues (Figure 2B). Broadening of the signals indicates that the metal-bound IRT1 loop undergoes dynamic interchange among different metal-bound states. Interestingly, the mutant peptides with only the two first (H162 and H164) or two last histidine residues (H166 and H168) from the stretch mutated to uncharged residues were still able to bind Zn^2+^ (Figure S3). Furthermore, the signal for the D173N mutant peptide no longer responded to the addition of Zn^2+^ (Figure 2C, F), suggesting that D173 is indeed involved in metal coordination.

Finally, we analyzed the chemical shift variations in the absence and presence of 2 equivalents Zn^2+^ of each residue. Here, we found that the residues located at the N-terminal side of the histidine-rich fragment were very little affected by Zn^2+^ binding, whereas those located on the C-terminus underwent significant chemical shift variations (Figure S4A). The same behavior was observed for peptides with mutations of two histidine residues and for the aspartic acid mutant (Figure S4B, C, D), although to a lesser extent than for the wild-type peptide. As expected, the peptide with the four histidine residues mutated presented no chemical shift variation (Figure S4E).

These results are in line with the amino acid composition of IRT1 loop, which is rich in negatively charged amino acids (E and D) on its C-terminal side with an isoelectric point of 3.84 (a.a. 168-185), instead of 5.83 for its N-terminal side (a.a. 144-167).

### Metal binding affinities of IRT1 regulatory loop

In order to quantify more precisely the interaction between the regulatory loop of IRT1 and non-iron metals, we performed microscale thermophoresis (MST) experiments. We first used Zn^2+^ as substrate, as we previously showed that it binds to the IRT1 peptide by ICP-MS (24), MST (16) and by CD (this study), and Zn^2+^ excess was reported to drive IRT1 endocytosis (24). A constant concentration of the peptide was titrated with a one in half-serial dilution of the ligand, ZnCl_2_, and thermophoresis was measured. A binding affinity of 33 ± 2.1 nM, when fitted to the K_D_ model, was obtained for wild-type IRT1 loop assuming a 1:1 stoichiometry of the reaction (Figure 3A). Data points from the peptide mutated for the four histidine residues (4HA) were recorded and failed to fit a binding curve, confirming the absolute requirement of the histidine stretch for metal binding (Figure 3A) (16). To evaluate deeper the contribution of the residues from the histidine stretch, we decided to test double mutant peptides harboring double H162A/H164A or H166A/H168A mutations. Interestingly, both double histidine substitutions resulted in reduced affinities for Zn^2+^ compared to WT, with a K_D_ of 240 ± 73 nM for the H162A/H164A mutant and a K_D_ of 234 ± 86 nM for the H166A/H168A mutant (Figure 3A).

**Figure 3.**
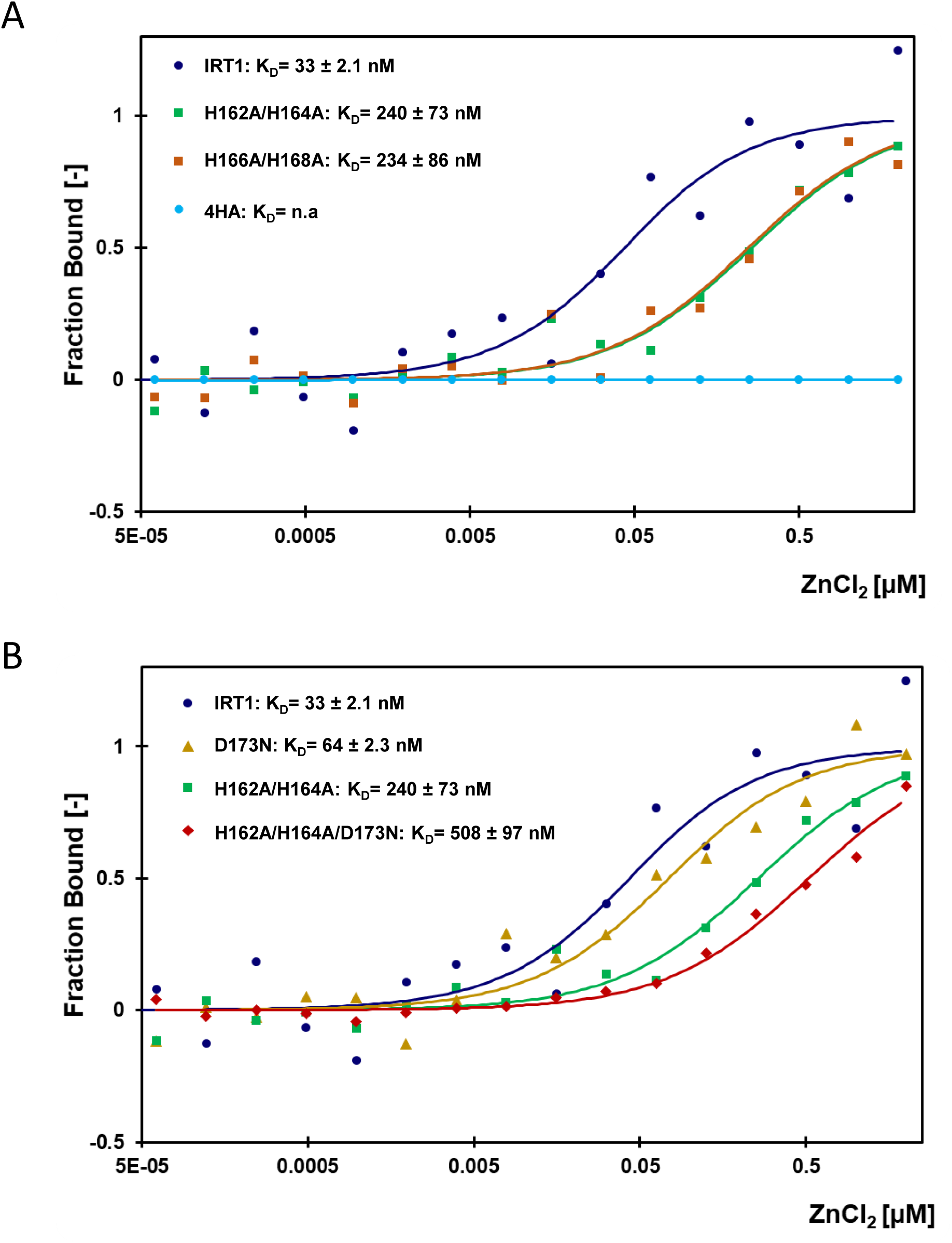
Zinc binding affinities of IRT1 regulatory loop. A) Microscale thermophoresis analyses of zinc binding to wild-type IRT1 (144-185) (IRT1; dark blue), double mutant with histidine residues 162 and 164 mutated to alanine (H162A/H164A; green), double mutant with histidine residues 166 and 168 mutated to alanine (H166A/H168A; orange) and quadruple mutant with histidine residues 162, 164, 166 and 168 mutated to alanine (4HA; light blue). Dots represent the average dose response of biological replicates. Errors bars and MST binding parameters are shown in Figure S8. B) Microscale thermophoresis analyses of zinc binding to wild-type IRT1 (144-185) (IRT1; dark blue), single mutant with aspartic acid 173 mutated to asparagine (D173N; yellow), double mutant with histidine residues H162 and H164 mutated to alanine (H162A/H164A; green), and triple mutant with histidine residues 162 and 164 mutated to alanine and aspartic acid 173 mutated to asparagine (H166A/H168A/D173N; red). Dots represent the average dose response of biological replicates. Errors bars and MST binding parameters are shown in Figure S8.

To inquire about the role of the D173 residue in metal binding affinity, the IRT1 loop carrying the D173N mutation was subjected it to MST analyses. Binding curves for Zn^2+^ were determined in the same concentration range as the previous experiments and fitted to the K_D_ model. The binding curve obtained for the D173N mutant was slightly shifted to the right compared to the one obtained with the wild-type peptide (Figure 3B), with a K_D_ of 64 ± 2.3 nM (Figure 3B). Because the contribution of D173 is likely minor compared to histidine residues, we sought to investigate the contribution of D173 in a peptide carrying a double histidine mutation where Zn^2+^ coordination is already destabilized, as shown above. The peptide carrying the three mutations showed significantly lower affinity to Zn^2+^ compared to the peptide with only two histidine residues mutated (Figure 3B). Such decrease in the affinity due to the removal of the negative charge from D173, in the context of the H162A/H164A mutation, argues for a contribution of D173 to Zn^2+^ coordination during sensing by the IRT1 transceptor. Considering that IRT1 also transports Mn^2+^ and that this non-iron metal was also shown to regulate IRT1 degradation (8,10,11,14,15,24), we monitored Mn^2+^ binding to IRT1 loop by MST. We observed that IRT1 loop is also able to bind Mn^2+^ at nanomolar range, with lower affinity than for Zn^2+^, showing a K_D_ of 114 ± 62 nM (Figure 4A). Similar to what has been observed with Zn^2+^, the four histidine IRT1 mutant loop was no longer able to bind Mn^2+^ and the double histidine mutants presented lower affinity compared to wild-type IRT1. Interestingly, the absence of two histidine residues showed less impact on the affinity for Mn^2+^ (Figure 4A) compared to Zn^2+^ (Figure 3A), pointing to the possibility of others residues being actively involved in Mn^2+^ coordination. To test if D173 could also participate to Mn^2+^ binding, we studied the affinity of the mutants harboring D173N mutation. Single D173N mutation barely affected affinity for Mn^2+^ compared to wild-type (Figure 4B). Nevertheless, D173N combined with H162A/H164A mutations resulted in a strong decrease for Mn^2+^ affinity (Figure 4B), revealing a strong contribution of these three residues also to Mn^2+^ coordination.

**Figure 4.**
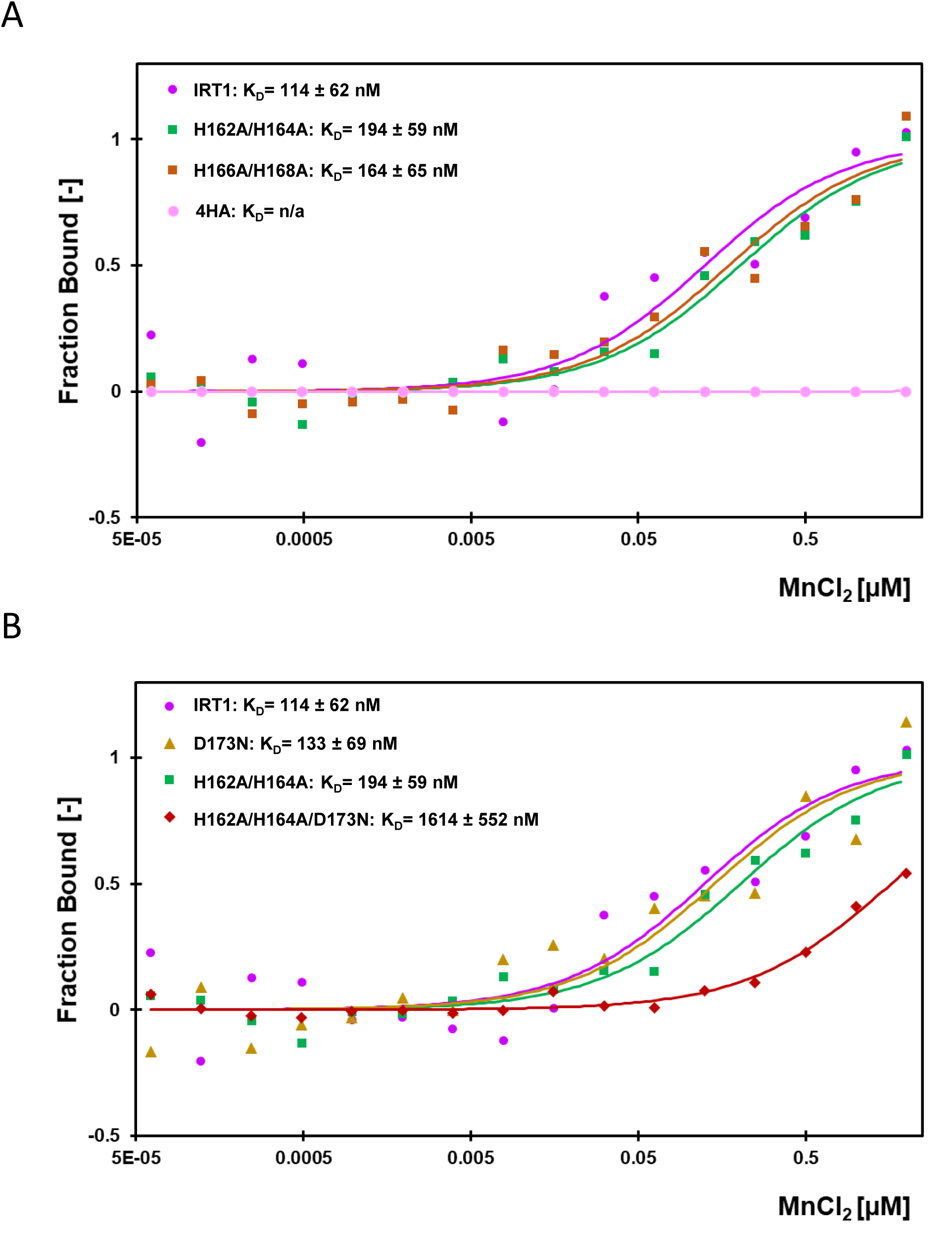
Manganese binding affinities of IRT1 regulatory loop. A) Microscale thermophoresis analyses of manganese binding to wild-type IRT1 (144-185) (IRT1; dark pink), double mutant with histidine residues 162 and 164 mutated to alanine (H162A/H164A; green), double mutant with histidine residues 166 and 168 mutated to alanine (H166A/H168A; orange) and quadruple mutant with histidine residues 162, 164, 166 and 168 mutated to alanine (4HA; light pink). Dots represent the average dose response of biological replicates. Errors bars and MST binding parameters are shown in Figure S9. B) Microscale thermophoresis analyses of manganese binding to wild-type IRT1 (144-185) (IRT1; dark pink), single mutant with aspartic acid 173 mutated to asparagine (D173N; yellow), double mutant with histidine residues H162 and H164 mutated to alanine (H162A/H164A; green), and triple mutant with histidine residues 162 and 164 mutated to alanine and aspartic acid 173 mutated to asparagine (H162A/H164A/D173N; red). Dots represent the average dose response of biological replicates. Errors bars and MST binding parameters are shown in Figure S9.

### Analysis of the role of aspartic acid D173 in planta

To characterize the role of residue D173 in metal transport and sensing, we first decided to express full length IRT1_D173N_ in the iron uptake-defective *fet3fet4* yeast mutant to evaluate if the corresponding protein is still active for iron transport. Yeast expressing wild-type IRT1 were able to complement the growth defect of *fet3fet4* in low iron conditions, as previously reported (10). Expression of IRT1_D173N_ protein yielded comparable growth compared to wild-type IRT1, indicating that IRT1_D173N_ is fully functional for iron transport (Figure S5).

An excess of non-iron metal substrates of IRT1 (Zn^2+^, Mn^2+^) leads to IRT1 depletion from plasma membrane and increases IRT1 accumulation in endosomes for later degradation by lytic vacuole (24). This safety mechanism to limit heavy metal toxicity in plants due to metal overaccumulation through IRT1 involves metal sensing by the histidine-rich motif located in the regulatory loop of IRT1 (16,24). Considering that the combination of D173N and H162A/H164A mutations impairs IRT1 loop affinity for Zn^2+^ and Mn^2+^ to a greater extent than the H162A/H164A mutation (Figures 3B and 4B), we wondered if this aspartic acid also participates to metal sensing and metal excess-dependent degradation of IRT1 protein. To test this hypothesis, we investigated the functional impact of the sole D173 mutation on IRT1 degradation in response to non-iron metal excess *in planta*. For this purpose, we took advantage of the previously reported fluorescently tagged version of IRT1, IRT1-mCitrine, which has been demonstrated to complement the *irt1-1* knockout Arabidopsis plants and *fet3fet4* (24). IRT1-mCitrine variants were transiently expressed in *Nicotiana benthamiana* leaves under the control of the 35S constitutive promoter to study IRT1 localization in response to high levels of non-iron metals. We however observed that the D173N substitution yielded retention of IRT1 into intracellular structures resembling the endoplasmic reticulum (ER). This is likely explained by the unexpected generation of an N-glycosylation site at the corresponding position (Figure S6B). We therefore substituted the D173 residue for glutamine (Q), which showed no impact on IRT1 protein localization at the plasma membrane under standard growth conditions (Figure S6A). IRT1-mCitrine was also observed mostly at the plasma membrane when expressed in *N. benthamiana* in the absence of non-iron metals (-/-; Figure 5A), similarly to what has been reported in stable Arabidopsis transgenic plants (15,16,24,29). A 3-hour treatment with non-iron metal excess (-/+++) led to IRT1-mCitrine depletion from plasma membrane and internalization in endosomes (Figure 5A, B), as already observed in roots (24).

**Figure 5.**
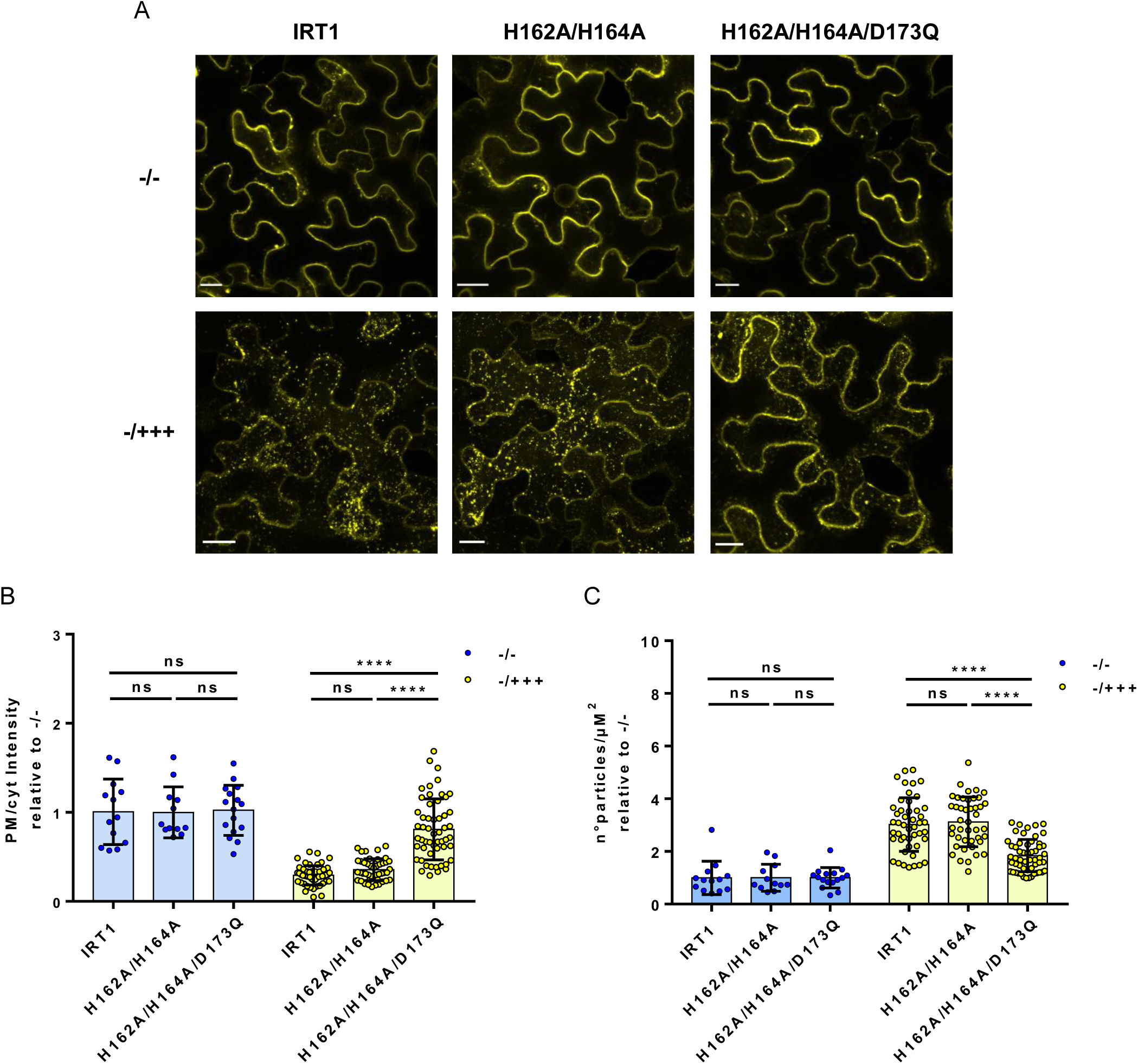
Residue D173 is involved in IRT1 endocytosis in response to non-iron metal excess. A) Representative confocal microscopy images of epidermal cells from *Nicotiana benthamiana* leaves transiently expressing 35::IRT1-mCitrine (IRT1) or mutated versions 35S::IRT1_H162A/H164A_-mCitrine (H162A/H164A) and 35S::IRT1_H162A/H164A/D173Q_-mCitrine (H162A/H164A/D173Q) after 3 hours of control treatment (without iron and non-iron metals; -/-) or after non-iron metal excess treatment (-/+++). Showed maximum projection of 10-15 optical sections taken using 1 µM z-distance. Scale bars, 20 µm. (B) Quantification of the ratio of the plasma membrane to intracellular signals and (C) quantification of intracellular particles per µM^2^ from cells exposed to metal excess relative to cells exposed to control solution of plants treated as described in A). Error bars represent SD (n=15-50) from 3 independent experiments. Asterisks indicate significant differences (one-way ANOVA, Tukey post-test, **** p < 0.0001).

This clearly indicates that *N. benthamiana* leaves possess the cellular machinery responsible for the internalization and degradation of IRT1 under metal excess. The IRT1 variant harboring the H162A/H164A double mutation exposed to a non-iron metal excess also displayed reduced plasma membrane localization (Figure 5A, B) and an increased number of endosomes (Figure 5C). Interestingly, the triple mutant H162A/H164A/D173Q showed significantly less protein removal from the plasma membrane (Figure 5A, B) and a significantly lower number of endosomes compared to double mutant H162A/H164A after treatment with high non-iron metal levels (Figure 5C). These observations argue for an implication of D173 residue in Zn^2+^ and Mn^2+^ sensing and proper IRT1 degradation when plants are experiencing non-iron metal stress.

## Discussion

To cope with changing nutrient availability and nutrient demand, plants developed strategies that allow them to control the expression of nutrient transporters. This is the case for the Arabidopsis IRT1 iron transporter that is controlled by various metal substrates at different levels, i.e. transcriptional regulation by low Fe^2+^ and post-translational regulation by Zn^2+^ and Mn^2+^ excess (24,29,30). Previously, we have shown that the post-translational regulation of IRT1 at the plasma membrane in response to non-iron metal excess involves its phosphorylation by the CIPK23 kinase followed by decoration with K63 polyubiquitin chains by the IDF1 E3 ligase, mechanism that require the histidine-rich motif in IRT1 cytoplasmic loop (24).

To gain further insight into the molecular mechanisms of metal sensing by IRT1, we here investigate the structural basis of the IRT1 regulatory loop using a combination of NMR spectroscopy, CD spectroscopy, MST analyses and protein prediction algorithms. We show that IRT1 loop is an intrinsically disordered region (IDR), which can adopt various conformations. This characteristic appears to be conserved among ZIPs, as it is also observed for the human ZIP4 zinc transporter (26,27). This is however not a specific feature of ZIPs since other families of transporters, like the Arabidopsis MTP1 vacuolar zinc transporter, also possess disordered intracellular loops with metal binding motifs (31). Our observations also indicate that IRT1 loop undergoes small structural changes in the presence of Zn^2+^, although it remains largely disordered. This observation points to the formation of a “fuzzy” complex between an IDR and a small ligand, as previously reported for hZIP4 (27). The enhanced flexibility and conformational plasticity of disordered regions in proteins provide a platform for post-translational events allowing enhanced interactions and chemical reactions between partners (32,33). For IRT1, such capacity may facilitate the recruitment of downstream factors such as the CIPK23 kinase for IRT1 loop phosphorylation upon non-iron metal excess (24). Moreover, the importance of the histidine stretch of the IRT1 regulatory loop in its post-translational regulation has been previously established, where plants harboring a protein mutated for the four histidine residues fail to show IRT1 phosphorylation and degradation upon non-iron metal excess (24). Interestingly, we observed chemical shift variations in the residues located at the C-terminus of the regulatory loop in the presence of Zn^2+^ compared to absence of Zn^2+^, region where the IRT1 residue predicted to be phosphorylated by CIPK23 kinase is found (24). Nevertheless, because the histidine resonances disappear upon the addition of Zn^2+^, we are not able to determine whether this particular stretch adopts a specific conformation in such conditions.

The nanomolar range dissociation constants recorded for Zn^2+^ and Mn^2+^ binding to IRT1 are consistent with experimentally determined nanomolar concentrations of such metals in plants grown in excess conditions (34). Importantly, we unambiguously demonstrate that the histidine stretch is of absolute importance for direct Zn^2+^ and Mn^2+^ coordination, as MST experiments demonstrated intense perturbations upon mutation of all four histidines, completely abolishing metal binding. Surprisingly, we uncovered that IRT1 loop is still able to bind Zn^2+^ when two histidine residues are mutated, probably due to the other two histidines and aspartic acid 173 being responsible for coordination. This is supported by the NMR data obtained with the double histidine mutants where proton resonance of the remaining two histidine residues and aspartic acid 173 still disappeared in presence of Zn^2+^. Although a coordination number of three for Zn^2+^ is not common, Zn^2+^ is described to adopt a variety of distorted coordination geometries without significant energy penalty (35). Besides, we speculate that IRT1 loop flexibility, as an IDR, could explain the remaining ability of triple mutant H164A/H166A/D173N to bind Zn^2+^, by allowing surrounding residues contribute to Zn^2+^ coordination, as predicted by Aphafold server 3 (36) for aspartic acid residue 144 (Figure S7).

Based on our NMR and MST observations, five residues from the regulatory loop of IRT1 could coordinate Zn^2+^. Since two Zn^2+^ ions have already been described to be coordinated by five ligands in proteins with one of the ligands would be acting as a bridge (37), two Zn^2+^-binding sites may also exist in IRT1 loop. This hypothesis is also supported by prediction made using Alphafold server 3 for IRT1 loop in presence of two Zn^2+^ ions, where histidine 168 is presented as the bridge coordinating both ions (Figure 6A). Unfortunately, the possible existence of two Zn^2+^-binding sites in IRT1 loop cannot be distinguished with our data. As postulated for the histidine-rich cytosolic loop of hZIP4 (27), we claim that a single Zn^2+^-bound state probably does not exist, but rather the Zn^2+^-bound state is likely a conformational ensemble with the Zn^2+^ coordinated by multiple combinations of the histidine residues and aspartic acid 173, allowing transient Zn^2+^ binding modes within IRT1 (144-185) depending on cytosolic Zn^2+^ concentrations. We also uncovered that the same residues involved in Zn^2+^ binding also participate to Mn^2+^ coordination, probably along with neighboring residues. We however demonstrate here that IRT1 loop shows higher affinity to Zn^2+^ compared to Mn^2+^, consistent with the stronger degradation of IRT1 observed in plants facings Zn excess (24). Metal ion availability and relative local cytoplasmic concentrations will likely dictate whether IRT1 loop will be coordinating Zn^2+^, Mn^2+^, or both.

**Figure 6.**
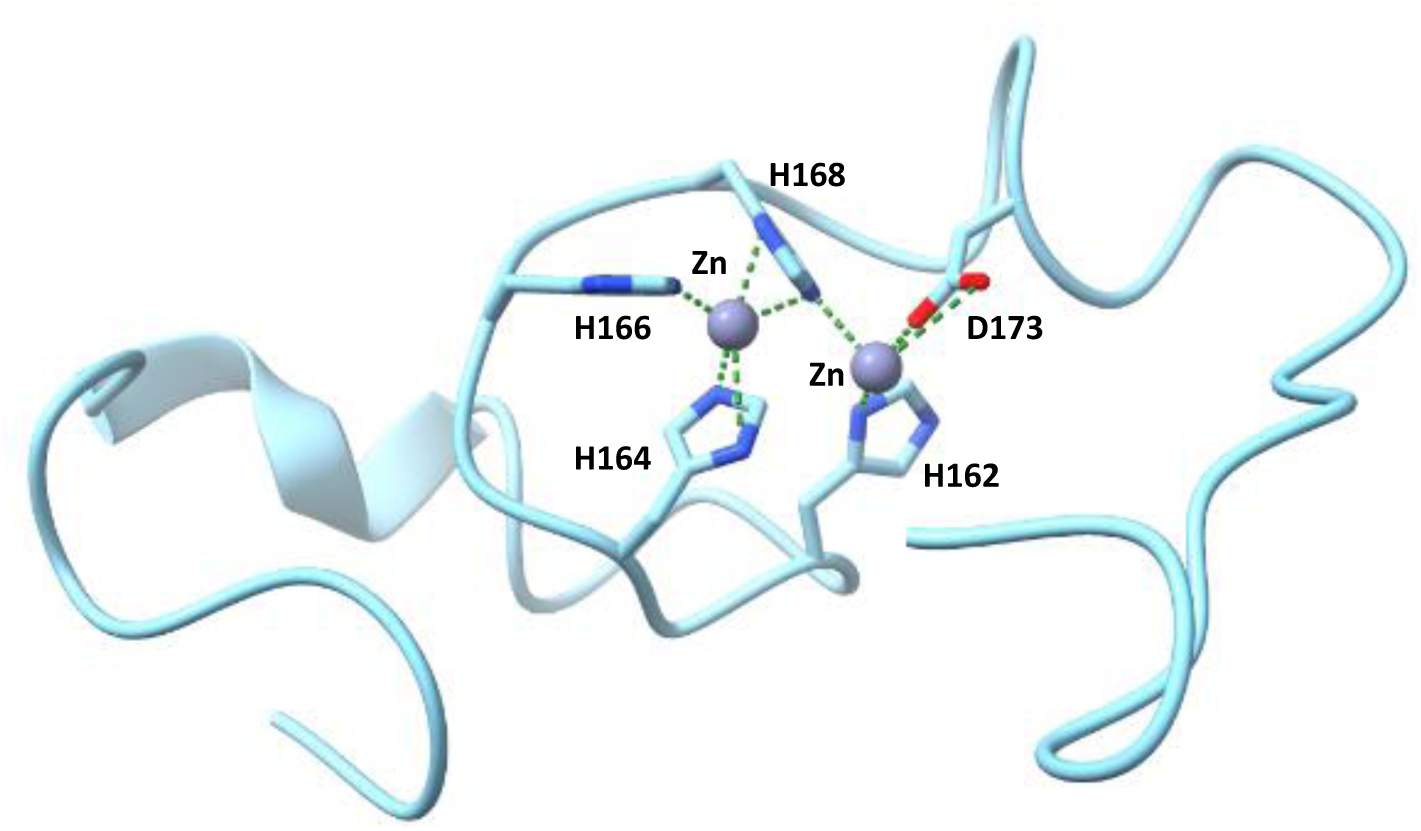
Zinc and manganese ion coordination by IRT1 regulatory loop. Alphafold3 prediction for IRT1 (144-185) coordinating Zn^2+^ ions (grey spheres) with residues H162, H164, H166, H168 and D173. Slashed green lines represent metal coordination within 3.5 Å distance. Analyses performed with USCF ChimeraX application.

Altogether, our work offers a framework for the analysis of metal sensing properties of ZIP transporters and deepens our understanding of how IRT1 protein senses metals through its regulatory loop. This understanding is crucial to grasp how plants optimize iron uptake and limit the absorption of highly-reactive non-iron metals in plant tissues and to consider biotechnological approaches to modulate heavy metal accumulation in plants.

## Experimental procedures

### Circular dichroism

Circular dichroism (CD) spectra were recorded on a Jasco J-815 spectropolarimeter equipped with a temperature controller operating at room temperature. CD spectra ranging from 190 to 240 nm were recorded in a 1-mm path length cuvette. Sample concentrations were at 25 µM, in 10 mM Tris, pH 7.0, and the data presented are an average of 3 scans.

### Sample preparation and NMR experiments

The wild-type IRT1 (144-185) peptide and the four mutant peptides were synthesized by Proteogenix with a purity grade of 95% minimum. Peptides were first dissolved at pH 3.5 to a final concentration of 500 µM in the presence of 150 mM NaCl. The pH was then adjusted to 6.7. Then, two equivalents of ZnCl_2_ for the peptide concentrations were added and the pH was again adjusted to 6.7. Two-dimensional phase-sensitive 1H Clean-TOCSY (38) with 60 ms spin lock, and NOESY experiments (38) with 200 ms mixing time was recorded at 5 and 20°C on an AVANCE Bruker 800.13 MHz spectrometer, with a spectral width of 8013 Hz, without sample spinning, with 2k real points in t2 and 512 t1-increments. Pulsed-field gradient-based WATERGATE was used for water suppression (39). The data were processed using TopSpin 4.0.6 software (Bruker). π/3 and π /6 phase-shifted sine bell window function was applied before Fourier transformation in both dimensions (t1 and t2). Data processing and analysis were performed using the Topspin ® 4.0 and CCPN NMR software (40).

### NMR structure of IRT1 (144-185)

Interproton distance restraints were derived from the two-dimensional ^1^H NOESY (with a 200 ms mixing time and at 5**°**C) using CcpNmr 2.4 (40) and used to generate IRT1 (144-185) structures with the program CYANA version 3.98.5 (41). We used the standard CYANA protocol of seven iterative cycles of NOE assignment and structure calculation, followed by a final structure calculation. In each cycle, the structure calculation started from 200 randomized conformers, and the standard CYANA simulated annealing schedule was used with 10,000 torsion angle dynamics steps per conformer. Graphic representations were prepared with PyMOL (42).

### Cloning expression and protein purification for Microscale thermophoresis

The wild-type IRT1 (a.a. 144-185) and mutant IRT1_4HA_ (a.a. 144-185) were cloned in the pMalc2x vector, in frame with the sequence encoding the Maltose Binding Protein (MBP) tag. H162A, H164A, H166A, H168A, and D173N mutations were introduced on the pMal-IRT1 (144-185) vector by primer reactions in parallel SPRINP method (43). Vectors were transformed into BL21 DE3 *E. coli* and recombinant proteins were purified using the manufacturer’s recommendation. Briefly, 1L of LB supplemented with glucose at 0.2% (w/v) final concentration and ampicillin was inoculated with 10 ml of an overnight high-density culture of cells containing the fusion plasmids pMal containing wild-type and mutants IRT1 (144-185). Cultures were grown at 37°C upon agitation until OD_600_ reached 0.5. Expression of the wild-type and mutants IRT1 (144-185) was induced by the addition of IPTG to a final concentration of 0.3 mM and cell cultures were incubated at 20°C overnight. Pellets were recovered by centrifugation at 10,000 x *g* for 40 minutes at 4°C, then suspended in HEPES 10 mM pH 7, NaCl 150 mM in 30 ml/L culture. Suspended cells were stored until further processing at −80°C.

Affinity purification of the MBP-IRT1 loop variants was performed in batches. Frozen cell suspensions were thawed. Lysozyme was added for cell wall disruption and then the suspensions were sonicated with 8 pulses for 45 seconds while kept on ice. Clarification of the lysate was performed at 20,000 x g for 30 min. The supernatant (crude extract) was recovered and stored in ice. Separately, 500 µL of amylose resin (NEB) was washed according to the provider’s specifications. Washed amylose resin was incubated with the crude extract in rotation at 4°C for 1h30. The resin was then pelleted at 500 g, and flow through was discarded. 2 washing steps were performed with HEPES 10 mM pH 7, NaCl 150 mM, 1 mM EDTA followed by two more washes in the same buffer without EDTA. Three consecutive elutions were carried out with HEPES 10 mM pH 7, NaCl 150 mM and Maltose 10 mM.

### Microscale thermophoresis

Binding experiments were performed by microscale thermophoresis with a Monolith NT.115 (NanoTemper® Technologies, Munich, Germany). MBP-IRT1 loop variants were labeled with the Monolith NT™ Protein Labeling Kit RED according to the instructions provided by the manufacturer, using a 1:3 protein:dye molar ratio. For binding experiments, the labeled proteins (20 nM) were incubated with a range of titrant concentrations made by serial dilutions (1:2), in 50 mM Tris buffer pH 7.4, 10 mM MgCl_2_, 150 mM NaCl, 0.05% Tween 20, in PCR tubes, at room temperature for 10 min. Premium treated capillaries (NanoTemper® Technologies) were loaded and the measurements were performed at 25 °C, 40 % LED power and 40 % microscale thermophoresis power, 20 s laser-on time and 1 s laser-off time. All the experiments were repeated at least twice with two independent protein labeling reactions. Binding data were analyzed using MO. AFFINITY ANALYSIS software (NanoTemper® Technologies).

### Yeast complementation assay

The complementation of *fet3fet4* yeast mutant was performed by expressing IRT1 or IRT1_D173N_ using the pDR195 yeast expression vector. Transformants were selected on a selective medium lacking uracil. For complementation assays, strains were grown at 28°C for 4 days on a selective medium without iron or containing 100 µM of Fe-EDTA.

### IRT1 constructs for plant expression

IRT1 mutations H162A, H164A and D173Q were introduced into the pDONR-IRT1mCitrine vector (24) using the SPRINP method (43). Final destination vectors were obtained by MultiSite Gateway® recombination using the entry vector described above, the pDONR-P4P1R-p35S entry vector containing the 2x35S promoter sequence, the pDONRP2RP3 entry vector containing a mock sequence, and the pGm43GW destination plasmid for expression in plants (24,44,45).

### IRT1 localization assays in Nicotiana benthamiana

*N.benthamiana* plants were grown for 4 weeks on soil under 16 h light at 22 °C and 55 % humidity before infiltration with *Agrobacterium tumefaciens* GV3101 strain.

IRT1-mCitrine variants were infiltrated into *A.tumefaciens* GV3110 and IRT1 protein localization assays were performed 48 h after agroinfiltration. Leaves transiently expressing IRT1-mCitrine variants were infiltrated with the treatments of study, consisting of control solution: half-strength Murashige and Skoog (MS/2) medium (46) lacking iron and non-iron metals; or metal excess solution: MS/2 lacking iron and containing Zn^2+^ (150 µM) and Mn^2+^ (300 µM). After 2 hours, leaves were re-infiltrated with correspondent treatment supplemented with 300 µM of cycloheximide for 1 h (in order to distinguish between newly IRT1 protein synthesized and endocytosed IRT1) and confocal images were taken.

### Confocal microscopy

For IRT1-mCitrine fluorescence in *N. benthamiana*, infiltrated leaves were mounted in discs in each correspondent treatment solution and confocal imaging was performed using a Leica TCS SP8 confocal laser scanning microscope (www.leica-microsystems.com/home/). For exciting mCitrine, the 514 nm laser line was used and emission was collected from 525 to 580nm. Laser intensity and detection settings were kept constant in individual sets of experiments.

### Data analyses of confocal images

Quantification of the ratio of the plasma membrane to cytosolic fluorescence signal and quantification of particles were calculated from the maximum projection of 10-15 optical sections taken using 1 µM z-distance. For the ratio, mean fluorescence was quantified using ImageJ software and results are presented as the mean value ± standard deviation (SD) of n=20-50 cell leaves from 3 independent experiments. Same cytoplasmic ROIs designed for measuring the ratio were used for quantifying particles using ComDet v.5.5.5 plugin from Image J software. One-way ANOVA was performed and the Tukey-post test method was applied to establish significant difference between means (p < 0.0001). Statistical analyses were performed using GraphPad Prism version 7.00 software.

### Bioinformatic tools

Alphafold tool (47) (https://alphafold.ebi.ac.uk/) and Alphafold3 server (https://golgi.sandbox.google.com) were used for IRT1 protein structure prediction; UCSF ChimeraX (https://www.rbvi.ucsf.edu/chimerax) for modeling IRT1 structure predictions and performing metal ion contact analyses; PONDR (48) (http://www.pondr.com) for predicting IRT1 degree of disorder and NetNGlyc (49) (https://services.healthtech.dtu.dk/service.php?NetNGlyc-1.0) was used for prediction of N-glycosylation sites in IRT1 protein.

## Data availability

All relevant data can be found within the manuscript and its supporting materials.

## Supporting information

This article contains supplementary information.

## Acknowledgments

We would like to acknowledge the Fédération de Recherche Agrobiosciences Interactions et Biodiversité of Toulouse (FRAIB) for the access to confocal microscopes and microscale thermophoresis systems.

