## Supplemental Figures 1-9 for "Metal sensing properties of the disordered loop from the Arabidopsis metal transceptor IRT1"

**Supplemental figure legends**

**Figure S1. Predicted structure and disorder regions of the IRT1 protein.**

A) Tridimensional structure of IRT1 predicted by Alphafold. The sequence of IRT1 (144-185) is shown. Disorder region is labeled in red.

B) Prediction of disordered regions obtained by PONDR®. Scores higher than 0.5 correspond to disordered regions and two arrows highlight IRT1 (144-185).

**Figure S2. Residues AVGI make a turn in IRT1 regulatory loop.**

A) Fingerprint region of the 1H-1H NOESY spectrum, showing the intramolecular and intermolecular NH-CHβ signals, recorded on IRT1 (144-185). The labels in black indicate the HN(i), H α (i) correlations and in red those corresponding to the HN(i), H α (i-2), and HN(i), H α (i-3) of amino acid I159.

B) Superimposition of IRT1 (144-185) structures calculated by the DYANA program.

C) Structure of the turn involving the three hydrophobic residues A156, V157 and I159 in licorice representation.

**Figure S3. Zinc binds to histidine residues in the IRT1 regulatory loop.**

Fingerprint region of the $^{1}$H-1H NOESY spectrum, recorded on the indicated two histidines double mutants of IRT1 (144-185) in the presence or absence of $Zn^{2+}$, showing several intramolecular and intermolecular NH-CHβ signals (A and B) and intramolecular CHα-CHβ signals (C and D).

The sequences of the domain into which mutations have been introduced are shown above each spectrum and the mutations are highlighted in green. The spectra recorded in the absence of

$Zn^{2+}$ are shown in black and those recorded in the presence of two molar equivalents of $Zn^{2+}$ are shown in red. * Indicate impurities in the sample. Annotations with a dash indicate cross-peak between inter residue i.e. NHi-CHβi-1, and without dash cross-peaks between intra residue i.e. HNi-Hβi and Hαi-Hβi.

**Figure S4. Chemical shift variations in the presence and absence of zinc.**

Chemical shift variations were determined for the wild-type (A) and indicated mutant peptides (B-E) in the absence or presence of two equivalents of $Zn^{2+}$. Sequences of the peptides are indicated at the top of each graph. Superimposition of chemical shift variations of the mutant peptides (red) and wild-type IRT1 peptide (black) is shown.

**Figure S5. $IRT1_{D173N}$ expression in yeast.**

Yeasts transformed with the pDR195 empty vector, or pDR195 carrying IRT1 or $IRT1_{D173N}$ were spotted on media without iron or on media with 100 μM of Fe-EDTA. Images were taken after 4 days.

**Figure S6: Impact of D173N and D173Q mutations on IRT1 localization.**

A) Confocal microscopy images of epidermal cells from *Nicotiana benthamiana* plants transiently expressing IRT1 mutated versions fused to mCitrine fluorescence tag under 35S promoter: 35S::$IRT1_{D173N}$-mCitrine and 35S::$IRT1_{D173Q}$-mCitrine. IRT1 protein harboring asparagine (N) residue in the 173 position retains IRT1 at the endoplasmic reticulum while glutamine (Q) mutation allows proper IRT1 localization at the plasma membrane. Scale bars, 20 μm.

B) N-glycosylation sites predicted for $IRT1_{D173N}$ and $IRT1_{D173Q}$ by NetNGlyc-1.0 server. The N residue at position 173 of IRT1 is predicted to be highly glycosylated, which could explain IRT1 retention in the endoplasmic reticulum, unlike the Q residue.

**Figure S7: Zinc coordination by triple mutated IRT1 regulatory loop.**

Alphafold3 prediction for IRT1(144-185) triple mutant H162A/H164A/D173N coordinating $Zn^{2+}$ ion (grey sphere) with residues H166, H168 and D144. Slashed green lines represent metal coordination within 3.5 Å distance. Analyses performed with USCF ChimeraX

**Figure S8: Microscale thermophoresis data for $Zn^{2+}$ binding.**

Microscale thermophoresis analyses of zinc binding by wild-type IRT1 (144-185) (IRT1; dark blue), single mutant with aspartic acid 173 mutated to asparagine (D173N; yellow), double mutant with histidine residues 162 and 164 mutated to alanine (H162A/H164A; green), double mutant with histidine residues 166 and 168 mutated to alanine (H166A/H168A; orange), triple mutant with histidine residues 162 and 164 mutated to alanine and aspartic acid 173 mutated to asparagine (H162A/H164A, D173N; red), and quadruple mutant with histidine residues 162, 164, 166 and 168 mutated to alanine (4HA; light blue). Dots represent the average dose response of biological replicates with errors bars. Table shows MST binding parameters comprising Standard Error of Regression, Signal to Noise and Response Amplitude values.

**Figure S9: Microscale thermophoresis data for $Mn^{2+}$ binding.**

Microscale thermophoresis analyses of manganese binding by wild-type IRT1 (144-185) (IRT1; dark pink), single mutant with aspartic acid 173 mutated to asparagine (D173N; yellow), double mutant with histidine residues 162 and 164 mutated to alanine (H162A/H164A; green)

double mutant with histidine residues 166 and 168 mutated to alanine (H166A/H168A; orange), triple mutant with histidine residues 162 and 164 mutated to alanine and aspartic acid 173 mutated to asparagine (H162A/H164A, D173N; red), and quadruple mutant with histidine residues 162, 164, 166 and 168 mutated to alanine (4HA; light pink). Dots represent the average dose response of biological replicates with errors bars. Table shows MST binding parameters comprising Standard Error of Regression, Signal to Noise and Response Amplitude values.

Fig S1

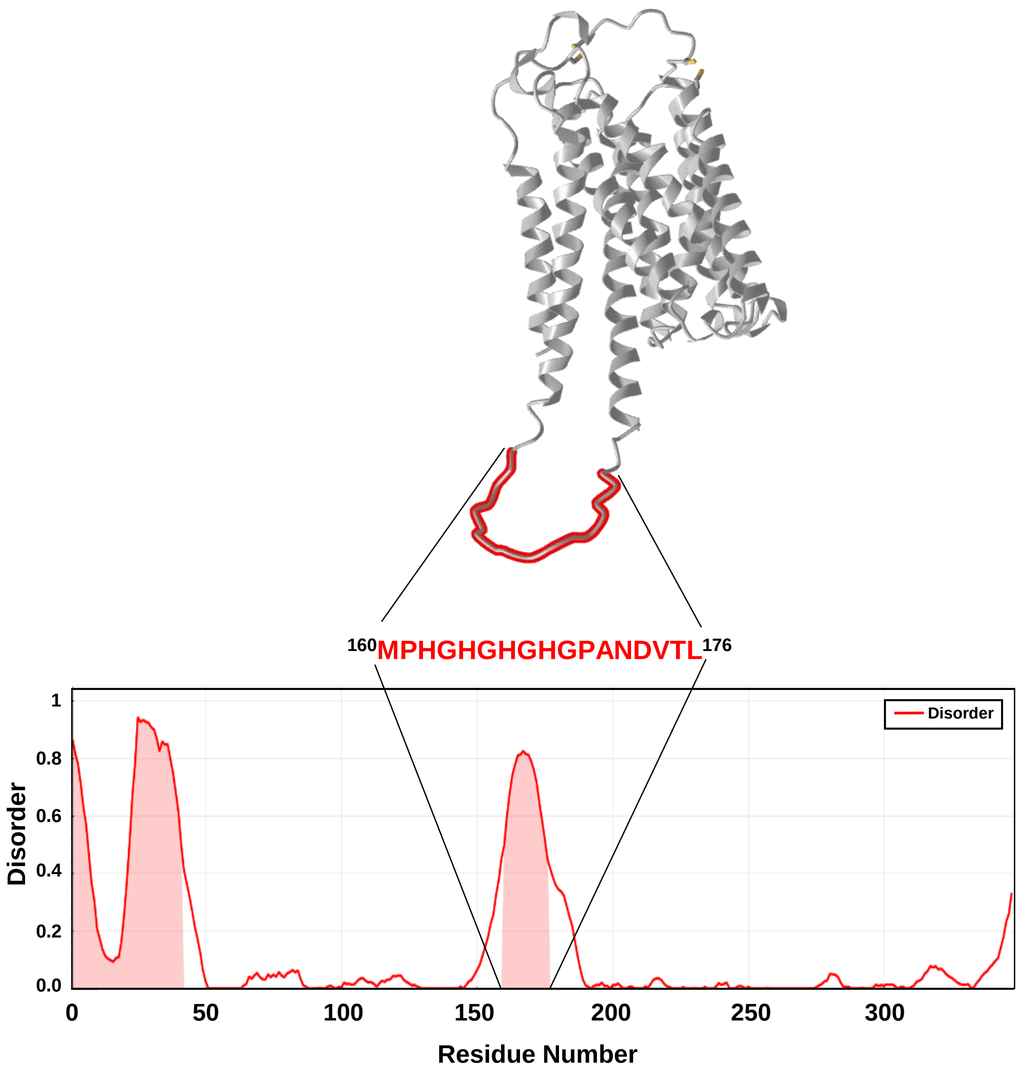
160MPHGHGHGHGPANDVTL176
Disorder
Disorder
1
0.8
0.6
0.4
0.2
0.0
0
50
100
150
200
250
300
Residue Number

Fig S2

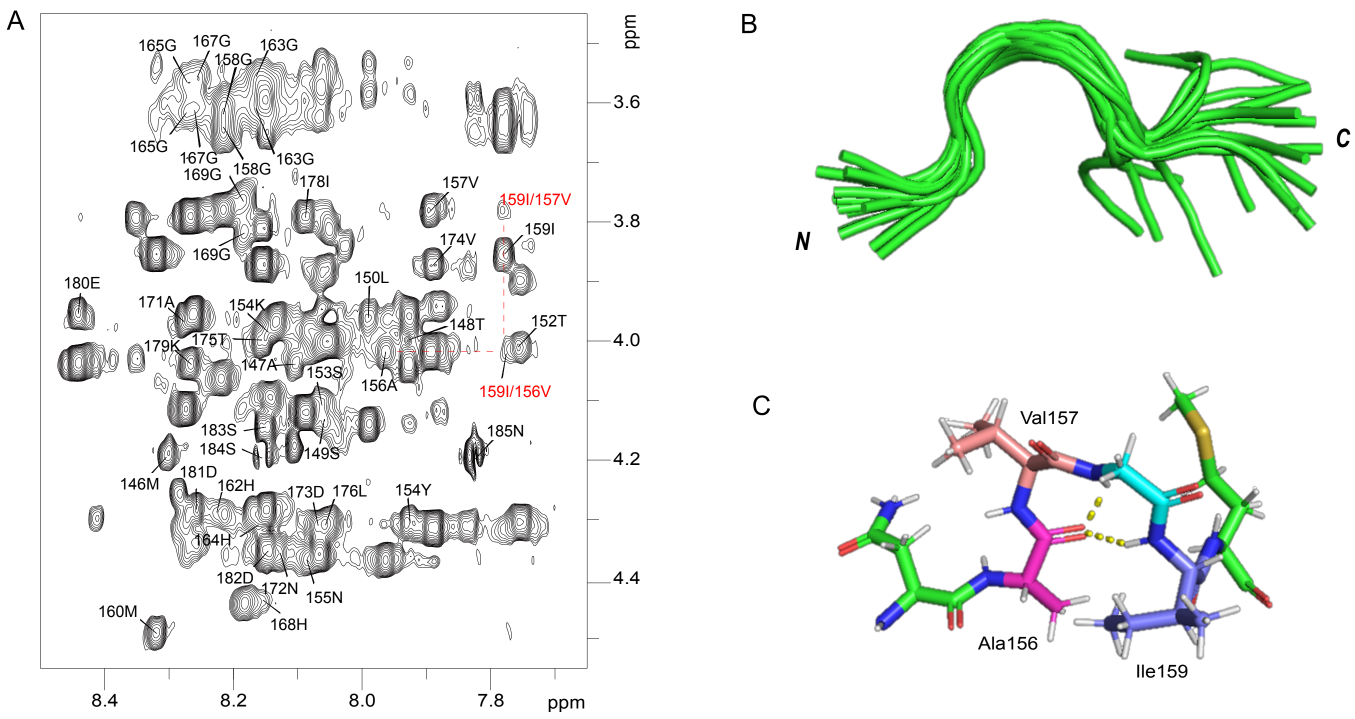

A
B
C
ppm
3.6
3.8
4.0
4.2
4.4
8.4
8.2
8.0
7.8
ppm
165G
167G
158G
163G
165G
167G
158G
163G
169G
178I
157V
159I/157V
159I
169G
174V
180E
150L
171A
154K
148T
152T
179K
175T
147A
153S
156A
159I/156V
183S
185N
184S
149S
146M
181D
162H
173D
176L
154Y
164H
182D
172N
155N
160M
168H
N
C
Val157
Ala156
Ile159

Fig S3

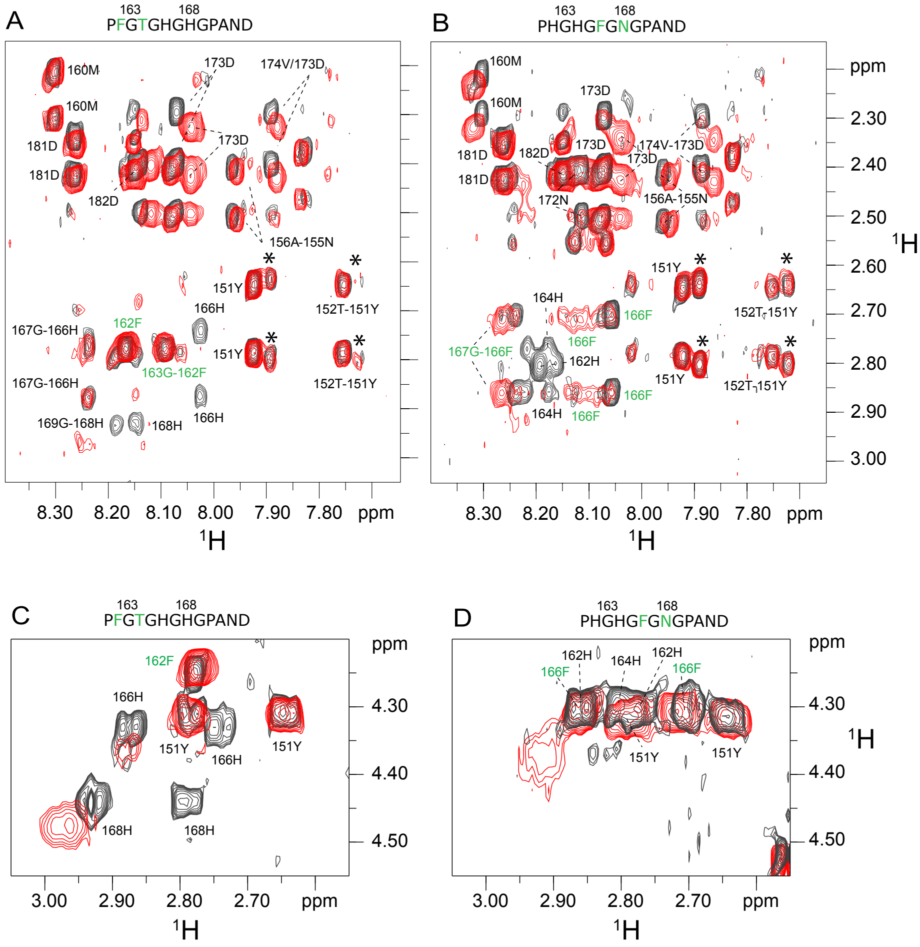

A
163 168
PFGTGHGHGPAND
160M
160M
181D
181D
182D
173D
174V/173D
173D
156A-155N
151Y
152T-151Y
166H
162F
167G-166H
151Y
163G-162F
152T-151Y
167G-166H
166H
169G-168H
168H
8.30 8.20 8.10 8.00 7.90 7.80 ppm
¹H
B
163 168
PHGHGFGNGPAND
160M
160M
173D
181D
182D
173D
174V-173D
173D
181D
172N
156A-155N
151Y
152T-151Y
164H
166F
166F
167G-166F
162H
151Y
152T-151Y
166F
164H
166F
8.30 8.20 8.10 8.00 7.90 7.80 ppm
¹H
ppm
2.30
2.40
2.50
2.60
2.70
2.80
2.90
3.00
¹H
C
163 168
PFGTGHGHGPAND
162F
166H
151Y
166H
151Y
168H
168H
ppm
4.30
4.40
4.50
3.00 2.90 2.80 2.70 ppm
¹H
D
163 168
PHGHGFGNGPAND
162H
164H
162H
166F
166F
151Y
151Y
ppm
4.30
4.40
4.50
¹H
3.00 2.90 2.80 2.70 ppm
¹H

Fig S4

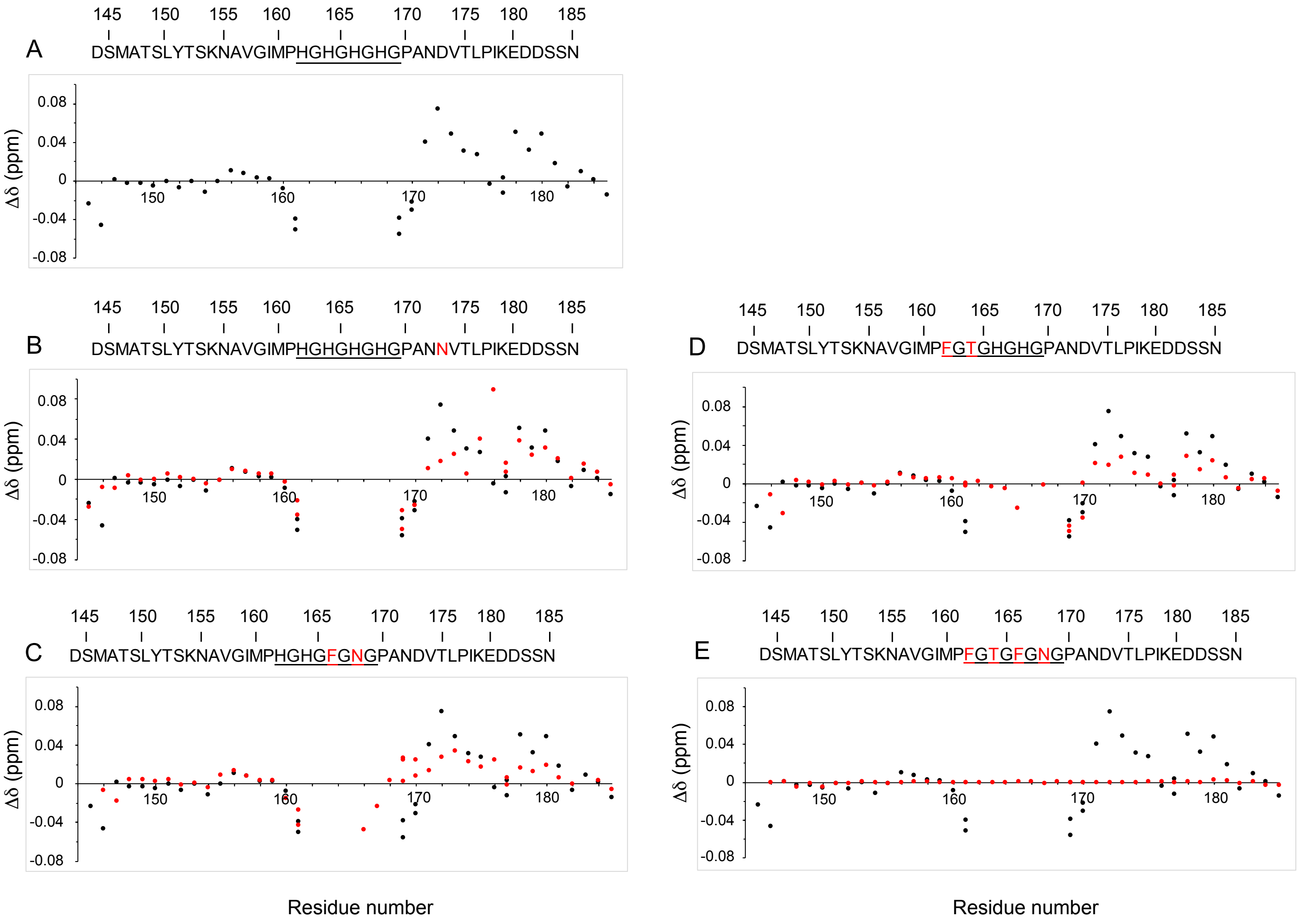
A
DSMATSLYTSKNAVGIMPHGHGHGHGPANDVTLPIKEDDSSN
B
DSMATSLYTSKNAVGIMPHGHGHGHGPANNVTLPIKEDDSSN
C
DSMATSLYTSKNAVGIMPHGHGFGNGPANDVTLPIKEDDSSN
D
DSMATSLYTSKNAVGIMPFGTGHGHGPANDVTLPIKEDDSSN
E
DSMATSLYTSKNAVGIMPFGTGFGNGPANDVTLPIKEDDSSN
145
150
155
160
165
170
175
180
185
Δδ (ppm)
0.08
0.04
0
-0.04
-0.08
Residue number
Residue number

Fig S5

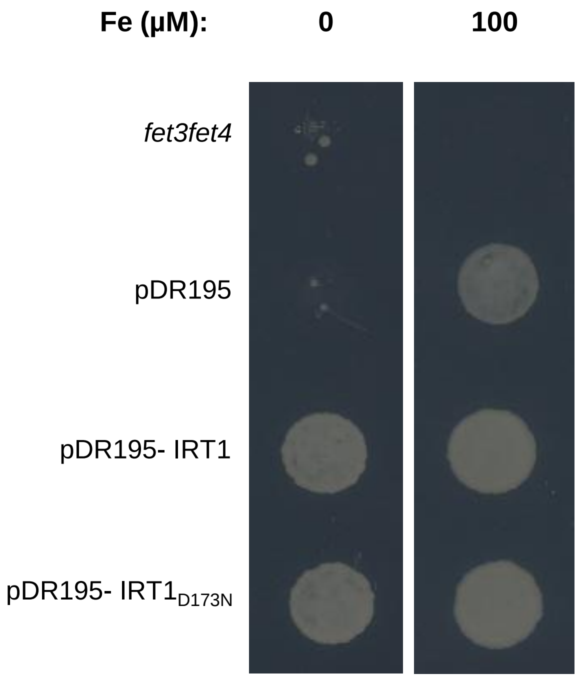

Fig S6

A

**D173N**

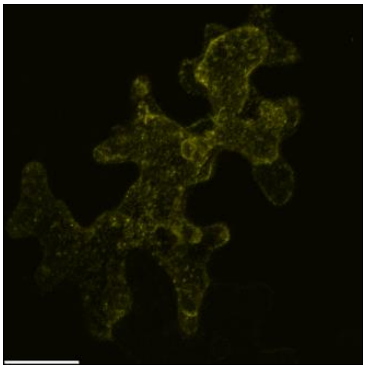

B

**D173N**

```
MASNSALLMKTIFLVLIFVSFAISPATSTAPEECGSESANPCVNKAKALPLKVIAIFVILIASMIGVGAPLFSRNVSFLQ      80
PDGNIFTIIKCFASGIILGTGFMHVLPDSFEMLSSICLEENPWHKFPFSGFLAMLSGLITLAIDSMATSLYTSKNAVGIM     160
PHGHGHGHGPANNVTLPIKEDDSSNAQLLRYRVIAMVLELGIIVHSVVIGLSLGATSDTCTIKGLIAALCFHQMFEGMGL     240
GGCILQAEYTNMKKFVMAFFFAVTTPFGIALGIALSTVYQDNSPKALITVGLLNACSAGLLIYMALVDLLAAEFMGPKLQ     320
GSIKMQFKCLIAALLGCGGMSIIAKWA
..........................................................................N.....      80
................................................................................     160
............N...................................................................     240
................................................................................     320
...........................                                                          400

(Threshold=0.5)
----------------------------------------------------------------------
SeqName      Position  Potential   Jury    N-Glyc
                                 agreement result
----------------------------------------------------------------------
D173N          75 NVSF   0.5248    (6/9)    +
D173N         173 NVTL   0.7439    (9/9)    ++
----------------------------------------------------------------------
```

**D173Q**

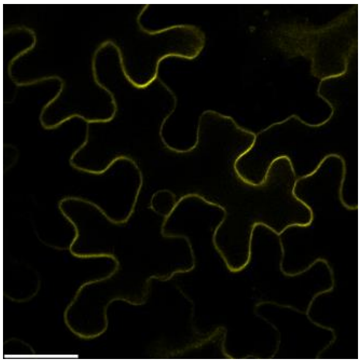

**D173Q**

```
MASNSALLMKTIFLVLIFVSFAISPATSTAPEECGSESANPCVNKAKALPLKVIAIFVILIASMIGVGAPLFSRNVSFLQ      80
PDGNIFTIIKCFASGIILGTGFMHVLPDSFEMLSSICLEENPWHKFPFSGFLAMLSGLITLAIDSMATSLYTSKNAVGIM     160
PHGHGHGHGPANQVTLPIKEDDSSNAQLLRYRVIAMVLELGIIVHSVVIGLSLGATSDTCTIKGLIAALCFHQMFEGMGL     240
GGCILQAEYTNMKKFVMAFFFAVTTPFGIALGIALSTVYQDNSPKALITVGLLNACSAGLLIYMALVDLLAAEFMGPKLQ     320
GSIKMQFKCLIAALLGCGGMSIIAKWA
..........................................................................N.....      80
................................................................................     160
................................................................................     240
................................................................................     320
...........................                                                          400

(Threshold=0.5)
----------------------------------------------------------------------
SeqName      Position  Potential   Jury    N-Glyc
                                 agreement result
----------------------------------------------------------------------
D173Q          75 NVSF   0.5249    (6/9)    +
----------------------------------------------------------------------
```

Fig S7

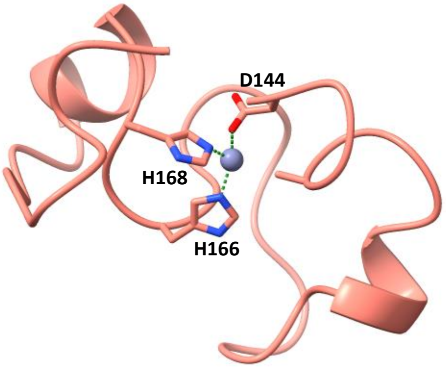

Fig S8

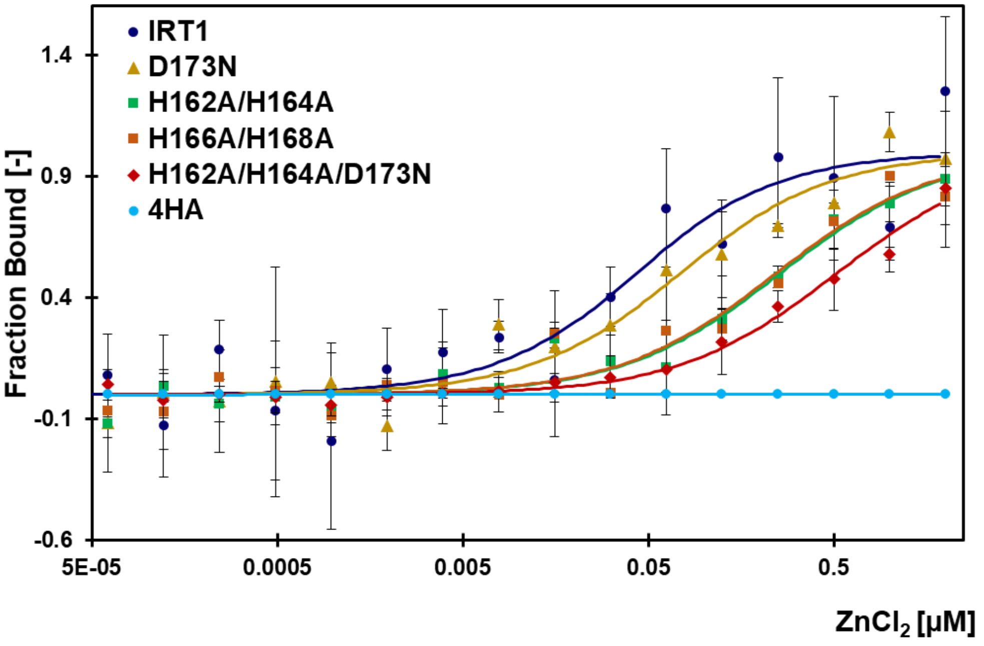


| Target | $K_D$ | SE of reg. | Sig. to noise | Resp. Amplitude |
|---|---|---|---|---|
| **IRT1** | 33 ± 2.1 nM | 0.38 | 6.3 | 2.25 |
| **H162A/H164A** | 240 ± 73 nM | 0.36 | 14.7 | 4.52 |
| **H166A/H168A** | 234 ± 86 nM | 0.40 | 12.2 | 4.57 |
| **4HA** | n.a | n.a | n.a | n.a |
| **D173N** | 64 ± 2.3 nM | 0.64 | 10.8 | 6.47 |
| **H162A/H164A D173N** | 508 ± 97 nM | 0.35 | 30 | 9.96 |

Fig S9

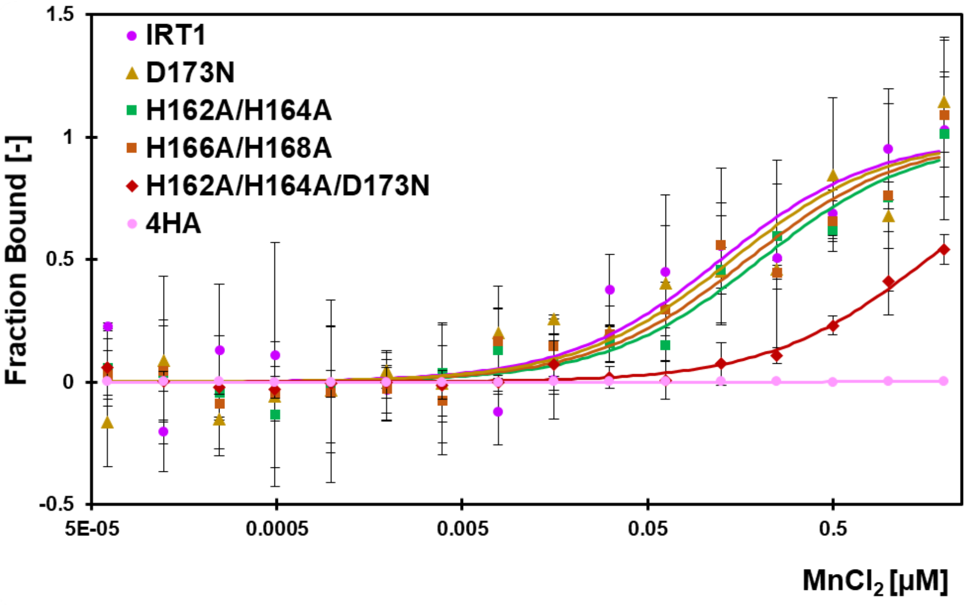


| Target | $K_D$ | SE of reg. | Sig. to noise | Resp. Amplitude |
|---|---|---|---|---|
| **IRT1** | 114 ± 6.2 nM | 0.29 | 6.3 | 1.99 |
| **H162A/H164A** | 194 ± 59 nM | 0.37 | 14.7 | 4.90 |
| **H166A/H168A** | 164 ± 65 nM | 0.64 | 12.2 | 6.30 |
| **4HA** | n.a | n.a | n.a | n.a |
| **D173N** | 133 ± 69 nM | 0.60 | 10.8 | 4.36 |
| **H162A/H164A D173N** | 1614 ± 552 nM | 0.35 | 30 | 11.81 |
